# Mobile Eye Tracking in the Real World: Best Practices

**DOI:** 10.1101/2025.08.11.669631

**Authors:** Debora Nolte, Jasmin L. Walter, Lane von Bassewitz, Jonas Scherer, Martin M. Müller, Peter König

**Affiliations:** Institute of Cognitive Science, University of Osnabrück, Osnabrück, Germany; Department of Neurobiology, University Bielefeld, Bielefeld, Germany; Department of Neurophysiology and Pathophysiology, University Medical Center Hamburg-Eppendorf, Hamburg, Germany

**Keywords:** eye tracking, mobile, GPS, spatial navigation, real world neuroscience

## Abstract

As research on human behavior, such as spatial navigation, increasingly adopts naturalistic settings, establishing best practices for such experiments becomes essential. While virtual reality (VR) offers a bridge between laboratory control and real-world complexity, it does not fully capture the experiential richness of real-world environments. Here, we present a demonstration of a mobile eye-tracking study conducted in a large-scale, outdoor urban environment, featuring unconstrained, long-duration free exploration and outside-pointing tasks. Using the city of Limassol, Cyprus as our testbed, we showcase the feasibility of collecting high-quality mobile eye-tracking, head orientation, and GPS data “in the wild,” capturing a wide range of natural behavior with minimal experimental constraints. Based on this experience, we provide a set of best practices tailored to the logistical and methodological challenges posed by complex, real-world urban settings, challenges unlikely to arise in traditional indoor or highly controlled environments. While these recommendations have general relevance, we exemplify them in the context of spatial navigation research. By establishing methodological standards for studies at this scale, we aim to encourage and inform future research into naturalistic human behavior outside the laboratory.

## 1 Introduction

Much of what we know about visual and cognitive behavior comes from highly controlled laboratory studies, which offer precision and reproducibility by using well-defined manipulations (Matusz et al., 2019; Vallet & van Wassenhove, 2023), typically involving seated participants, screen-based tasks, and minimal body movement (Matusz et al., 2019; Shamay-Tsoory & Mendelsohn, 2019). However, these experiments fail to capture the complexity and dynamics of natural behavior (Foulsham et al., 2011; Janssen et al., 2021; Land & Hayhoe, 2001; Nastase et al., 2020; Tatler et al., 2011; Vallet & van Wassenhove, 2023). This limitation is particularly apparent in domains such as spatial navigation, where interactions of body and environment play a central role (Chrastil & Warren, 2013; Taube et al., 2013; Wolbers & Wiener, 2014). As a result, in areas such as spatial navigation, laboratory-based studies may not provide the whole picture.

To address these limitations, immersive virtual reality (VR) has emerged as a promising approach. VR allows the study of cognition in rich, three-dimensional environments, enabling participants to move their eyes or bodies while maintaining experimental control and, to an extent, reproducibility of sensory input and task variables (Bohil et al., 2011; Slater & Sanchez-Vives, 2016; Thurley, 2022). VR allows participants to explore environments using naturalistic self-initiated movements and supports the integration of multisensory information (Diersch & Wolbers, 2019; Jeung et al., 2023; Thurley, 2022). For example, in spatial navigation research, VR-based studies have advanced our understanding of spatial knowledge acquisition (König et al., 2021), route learning (Hilton & Wiener, 2023), identification and use of landmarks (Walter et al., 2022), and the effects of social agents on navigation strategies (Sánchez Pacheco et al., 2025). Overall, VR provides a more ecologically valid context for studying embodied cognitive processes than lab-based setups while allowing for precise measurement and experimental control.

However, while VR has proven effective in capturing many aspects of real-world cognition, the question remains: to what extent do behaviors observed in VR translate to real-world environments? The absence of full-body motion, vestibular cues, or real-world unpredictability might alter how individuals perceive and interact with the environment in VR (Stangl et al., 2023). Although many studies report similarities (Hepperle & Wölfel, 2023; Pastel et al., 2022), others point to meaningful differences between the two contexts (Kalantari et al., 2024; Kimura et al., 2017). Therefore, while VR remains a powerful tool for investigating cognitive processes, it should be viewed as complementary to, rather than a substitute for, research conducted in natural, real-world environments.

Accordingly, research in real-world settings has gained increasing attention (Foulsham et al., 2011; Ladouce et al., 2017; Parada, 2018; Valtakari et al., 2021). Mobile eye-tracking and other wearable sensing technologies has enabled researchers to collect rich behavioral data in everyday environments (Ladouce et al., 2017; Makeig et al., 2009), making it possible to investigate cognitive processes as they naturally unfold in response to complex, real-world situations (Janssen et al., 2021; Land & Hayhoe, 2001; Nastase et al., 2020). Although these advances are promising, studies conducted outside the laboratory often face challenges such as unpredictable environmental factors, participant variability, data accuracy, and technical constraints, all of which can introduce noise and complicate replicability (Janssen et al., 2021; Matusz et al., 2019; Vigliocco et al., 2024). Furthermore, successfully implementing these studies involves a considerable learning curve, as researchers must navigate logistical, methodological, and technical challenges that differ significantly from those encountered in controlled laboratory environments. These factors highlight the pressing need for comprehensive methodological standards and practical guidelines on how to design, plan, conduct, and analyze high-quality cognitive experiments in real-world environments.

To work towards methodological standards and best practices for mobile real-world eye-tracking and spatial navigation research, we implemented a mobile eye tracking study and will discuss relevant aspects and considerations based on our experience and related literature. Specifically, we conducted a single-subject case study (28 years old, no history of neurological disorders, normal vision) replicating two established spatial navigation experiments in an immersive VR city (Sánchez Pacheco et al., 2025; Schmidt et al., 2023). Our participant freely explored the city center of Limassol, Cyprus, while we recorded eye movements, head orientation, and GPS data, followed by tasks assessing her spatial knowledge. As a co-author, the participant was informed about the study’s goals but was unaware of specific details of the experimental design. Using the example of this study, we outline methodological insights and concrete recommendations for researchers aiming to conduct (spatial navigation) studies in real-world settings with focus on mobile eye tracking, data stream synchronization, and general experiment settings.

## 2 Designing and Preparing the Experiment

Recording in the field, especially when far from the lab, requires sufficient planning and preparation before the recording begins. This may include a detailed plan of the experiment, practice of the experimental procedure, and adequate planning while at the location. In this work, we are specifically interested in visual exploration during spatial navigation. To this end, this project utilized mobile eye-tracking in a real-world city center as a participant performed spatial exploration and navigation tasks. In the following sections, we provide a comprehensive overview of essential aspects and best practices for designing a real-world experiment, using the specific example of our spatial navigation case study conducted in Cyprus.

### 2.1 Insights from VR Experiments

The design of this real-world study is adapted from prior work conducted in a large-scale VR city, “Westbrook” (Sánchez Pacheco et al., 2025; Schmidt et al., 2023), which served as a controlled environment for investigating spatial navigation and knowledge acquisition. Participants could freely explore this city while eye-tracking, body movements, and positional data were recorded. Following exploration, participants completed several navigation tasks to assess spatial knowledge. The results demonstrated, among others, that individual differences in spatial navigation performance can be explained with patterns in visual behavior during the spatial exploration (Walter et al., in preparation) and that human agents enhance visual exploration and improve spatial knowledge acquisition (Sánchez Pacheco et al., 2025). These rich VR datasets can also inspire different analysis approaches, including detailed examination of eye-tracking metrics (Clay et al., 2019; Nolte et al., 2024), and graph-theoretical analysis of viewing behavior during spatial knowledge acquisition (Walter et al., 2022). These studies show how eye-tracking data can be used to investigate spatial knowledge formation and provide motivation for experiments in the real world.

We recommend piloting novel experimental designs in controlled VR environments to inform about expected findings, analysis procedures, and potential results before moving to the real world.

### 2.2 Selecting the Experimental Area

Selecting an appropriate experimental area depends on the specific area of research and task requirements, which might vary drastically depending on the task demands. For instance, the size, shape, or landmarks within a selected area are all known to influence task difficulty and spatial navigation performance (Caduff & Timpf, 2008; Evans et al., 1984; Spiers et al., 2023). Therefore, it is important to carefully select an experimental area in a real-world environment based on the task demands and research question of interest.

For our spatial navigation study, we remotely pre-selected an experimental area. By using several online resources (Open Street Map, Google Maps, and Google Street View), we identified an area that matched the VR city Westbrook (Sánchez Pacheco et al., 2025; Schmidt et al., 2023) in size and the number and diversity of buildings, including unique facades, street art, and different building types like shops, cafes and churches. In addition, we tried to minimize salient global landmarks, such as the ocean shore and large roads. Ultimately, we selected an experimental area located in the Limassol city center (see Figure 1C). After physically exploring the experimental area in person and visualizing all walkable streets, our experimenter realized that the pre-selected area was too big for the current study and, consequently, decreased its size, spanning a total of 0.165 km² (see Figure 1C). The ocean and one big road were visible or audible from only a few points within the city, limiting their use as a global landmark, and were therefore considered acceptable. The area had a relatively consistent internet connection (see Section 2.8 *Piloting in the Real World* for remarks on this) and reliable GPS coverage (see Section 4.3 *GPS Data* for details), both of which were relevant for data collection.

**Figure 1:**
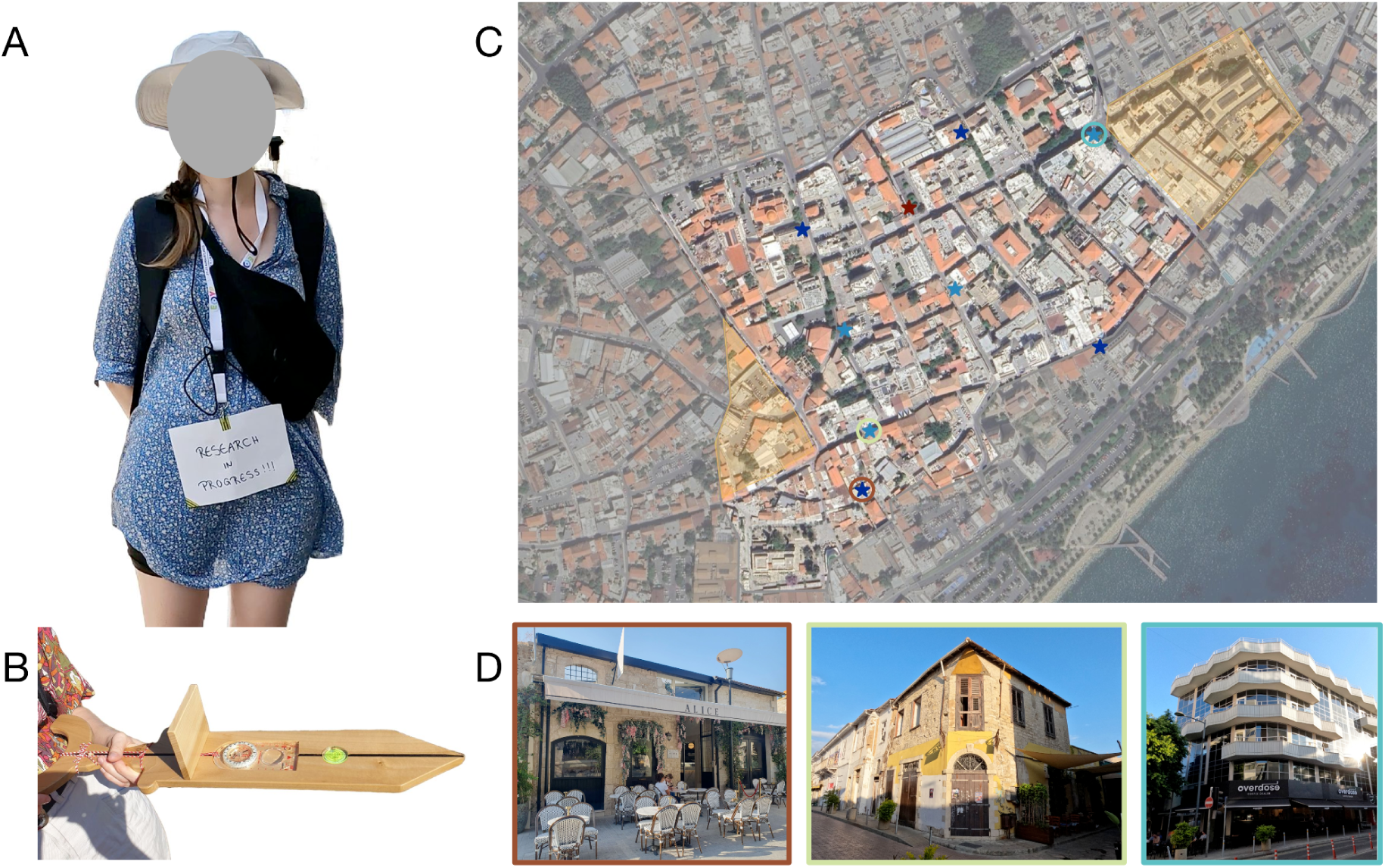
Designing the Experiment. (A) The participant was wearing Pupil Labs Neon eye-tracking glasses and a sunhat to improve data quality. The phone to record the eye-tracking data was stored in a cross-body bag. The GPS was attached to an antenna and carried in a backpack. (B) The sword used for the pointing tasks and to validate the head-tracker had an inbuilt compass and level. A removable sight protection prevented the participant from seeing the compass. (C) A map of the experimental area and task buildings. The selected experiment area is highlighted in bright colors, the excluded area is marked in grey. The parts of the city colored in yellow were excluded after an on-site visit by the experimenter. The red star indicates the starting location of the five continuous recording sessions. All task buildings are marked in blue, with dark blue indicating those that were also visited during the pointing tasks, while light blue indicates buildings that were only pointed to. The colorful circles around three task buildings correspond to the images of task buildings shown in (D). (D) Three examples of task buildings were shown; the pictures were identical to those presented to the participant during the pointing-to-building task. The borders around each image match the colorful circles displayed on the map.

The biggest takeaway from the real-world area was that it still felt too large for our purposes. One major factor was the participant’s walking speed, which was much lower than in VR (see Section 4.3 *GPS Data* for details). Navigation was further affected by moving elements such as pedestrians, vehicles, and varying weather or light conditions (see Section 2.7 *Effects of Weather and Sunlight* for details), changing and consequently making visual markers less reliable. The participant had to endure traffic, crowded spaces, and physical fatigue from walking in the heat, limiting her focus on the navigation task, and making it impossible to traverse all streets more than once. Moreover, the real-world environment contained a large number of buildings, many similar-looking facades, and narrow streets without gaps between buildings, and continued beyond the experimental area limits, thus adding additional visual stimuli beyond the experimentally relevant buildings. These factors highlight the need for a smaller experimental area or an adjustment of the spatial exploration or the experimental task.

We recommend carefully selecting the size, extent and composition of the area and verifying it on site by physically examining and experiencing the area before starting data collection. Having areas in pedestrian zones, wide pavements, good internet coverage, and the availability of detailed geospatial data and first-person video footage (i.e. digital urban twin, Open Street View, Google Street View) makes it easier to conduct and analyse experiments. The relevant aspects of selecting a good experimental area depend on the task, including the intended repetitions, and require careful and realistic consideration.

### 2.3 The General Experimental Design

#### 2.3.1 Spatial Exploration Phase

Designing spatial navigation tasks for real-world settings inspired by virtual ones (Sánchez Pacheco et al., 2025; Schmidt et al., 2023) may be straightforward in some parts but requires rethinking task structure, timing, and environmental compatibility for others. Real-world exploration must accommodate variable outdoor conditions, participant fatigue, and technical limitations, demanding a more flexible and resilient experimental design (Vigliocco et al., 2024).

Our study was structured into five sessions across three days, for a total exploration of 130 minutes. The first three sessions lasted 30 minutes each; the final two had to be spontaneously shortened to 20 minutes due to limited remaining time at the end of the day and our time in Cyprus, respectively. During each exploration session, the participant was freely able to explore the environment, familiarizing herself with the city’s layout and the buildings within. Each session began at a predefined central starting location (see Figure 1C) and was divided into ten-minute segments, providing regular breaks for both the humans and the equipment. As the participant remained within the experimental environment during these pauses, she was asked to keep her gaze lowered, or eyes closed, to minimize incidental exposure to spatial features. The equipment was calibrated and validated before and after each segment (see Section 3.2 *Preparing the Equipment* and Section 3.3 *Temporal Alignment of Data Streams* for details).

We recommend conducting short, timed sessions across multiple days, starting early enough to complete data collection by evening. Frequent, extended breaks are essential to prevent participant fatigue and accommodate unexpected technical issues, such as overheating of the equipment. Importantly, efforts should be made to prevent participants from being exposed to continued experimental information during these breaks.

#### 2.3.2 Spatial Navigation Tasks

Compared to lab-based studies, designing an experimental task in a real-world study requires planning to address logistical challenges and preserve data quality (Vallet & van Wassenhove, 2023). In particular, real-world task administration must contend with reduced control over timing, the lack of automated features available in digital simulations, and environmental constraints (Stangl et al., 2023; Vallet & van Wassenhove, 2023).

We assessed spatial knowledge using three tasks: a pointing-to-north task, a pointing-to-building task, and a map-drawing task. The pointing tasks were conducted via a mobile survey implemented with LimeSurvey that randomized pointing trial order, displayed images of preselected buildings, and stored participant responses. During preparation on site, we selected eight task buildings with a suitable area in front of them to serve as task locations, photographed them for the survey, and marked their locations on Google Maps (see 2.5 *Selecting the Task Buildings* for details). However, due to time constraints on site (see Section 3.7 *Time Management* for details), only four task locations were used during the task, although the participant still pointed to all eight original task buildings at these locations (see Figure 1C and 1D). In addition, the pointing task was performed at the starting location (see Figure 1C), directly before and after the pointing tasks trials. At each task location, the participant pointed north and then toward the remaining seven buildings in randomized order using a compass-equipped wooden sword (see Figure 1B). We recorded compass directions in the survey and logged participant’s locations using the RTK and manual markers in Google Maps for later angular error analysis (see Section 2.4 *The Equipment to Record the Data* and 5.5 *Task Performance* for details). After completing all trials, the participant was guided back to the starting location, where she performed a final round of pointing tasks and indicated her confidence (ranging from zero to ten) in her recall abilities. The experimenter noted down any comments she made regarding the buildings and her certainty in her performance. After completing all pointing tasks and after the sun had set, the participant drew her mental map of the city layout and relevant building locations on a tablet.

Real-world experiments require physically executing tasks and transferring digital or automated features to physical devices. Using a digital survey to randomize the pointing trials can be crucial to match task conditions to VR experiments and ensure a smooth task execution. Stationary tasks, such as map drawing, can extend data collection beyond daylight constraints.

### 2.4 The Equipment to Record the Data

Collecting data in real-world experiments poses a major challenge compared to VR, where information is automatically recorded in a controlled setting (Anderson et al., 2023; Clay et al., 2019). In outdoor environments, researchers must coordinate multiple devices, each with its own capabilities, limitations, and data format, while ensuring reliable performance across changing conditions and unpredictable external factors like signal loss and physical constraints (Shamay-Tsoory & Mendelsohn, 2019; Stangl et al., 2023; Vallet & van Wassenhove, 2023). However, with careful planning and consideration, successful data collection is possible.

For the purpose of our spatial navigation study, we recorded eye tracking data using the Pupil Labs Neon eye tracker (Baumann & Dierkes, 2023; see Figure 1A), which provided eye movements (sampling rate of 200 Hz and a resolution of 1600×1200 pixels), world camera footage (30 Hz RGB scene camera; 132° x 81° field of view; 1600×1200 pixels resolution), and head tracking data (IMU; recording accelerator, magnetometer, and gyroscope at 110 Hz). To record body position, we primarily used the Emlid Reach M+ RTK GNSS module (Emlid Tech, Budapest; see Figure 1A), which combines self-positioning via satellites (GPS, GALILEO, GLONASS, and Beidou) with real-time kinematic correction data from state or federal stationary reference points on the ground. The module provides positional data at centimeter-level precision when connected to such a correction reference point (Ng et al., 2018; Valente et al., 2020). Since this system relies on mobile network corrections, we collected backup GPS data using a GoPro Hero 10 (GoPro, Inc., San Mateo, CA) during exploration, and manually marked the participant’s positions in Google Maps during the pointing tasks. The GoPro’s primary function was to capture video footage of the participant exploring the city, specifically the participants’ direct surroundings and potential situations or people interfering with the data collection. Finally, to record the participant’s pointing direction during the pointing task, she used a custom-built wooden sword fitted with a digital compass and level (see Figure 1B). This device allowed us to accurately measure the pointing direction.

We recommend thoroughly testing all equipment in advance as well as on site to verify their function, and including backup systems wherever possible, as devices can be prone to failure in real-world settings. For collecting position data, a GoPro camera with GPS might suffice in place of the RTK system, depending on the required level of accuracy and the expected environmental conditions. If stationary, marking locations in Google Maps ahead of time and adjusting them if needed offers a practical alternative or sanity check. Overall, despite some challenges, we were satisfied with our setup and can recommend a similar configuration for future studies.

### 2.5 Selecting the Task Buildings

Transitioning from virtual to real-world experiments introduces specific challenges in selecting suitable experimental stimuli, such as choosing appropriate task buildings for a task assessing spatial knowledge via a pointing task. In virtual environments, researchers can manipulate the stimuli according to the experiment’s needs, for example, by distinguishing relevant buildings with graffiti (Sánchez Pacheco et al., 2025; Schmidt et al., 2023). In contrast, real-world studies must adapt to an existing environment, where similar distractors and uncontrolled visual conditions may affect attention and object recognition (Ladouce et al., 2019; Ringer et al., 2021). Moreover, individual differences in spatial abilities and navigation strategies influence which environmental features are used for orientation (Caduff & Timpf, 2008; Wolbers & Hegarty, 2010), potentially leading to variability in which buildings are remembered. Therefore, careful stimulus selection within a pre-existing environment is both crucial and challenging.

Our participant performed a spatial pointing task in an urban setting. The experimenter selected task-relevant buildings (see Figure 1D) matching the selection criteria in the VR experiments (Sánchez Pacheco et al., 2025; Schmidt et al., 2023) as well as adaptations emerging during the exploration phase on-site. Specifically, the task buildings had to be distributed equally across the experimental area and not visible from the corresponding task locations. Furthermore, matching the criteria in VR, the task buildings should contain buildings of varying size and function, while avoiding typical landmarks. Therefore, the selection of task buildings was updated several times on-site as the exploration progressed and in a last minute decision, considering the duration of the experiment, the task locations had to be reduced to ensure the completion of the tasks within the remaining time in Limassol. Nevertheless, despite the careful selection, the participant did not recognize all eight task buildings during the first trial of the pointing task. One of the unrecognized buildings also functioned as a task location. In response, the experimenter decided to use the task location of this unknown task building first. While this improved the participant’s performance and introduced meaningful variance in a low-sample context, it also resulted in a potential bias that could only be justified in the specific context of this experiment.

Stimulus selection in real-world studies is task and environment-dependent. We recommend defining clear selection criteria when choosing appropriate stimuli and validating these stimuli on-site. Considering fall-back options beforehand is important to help limit stressful decisions when under time pressure. When recording in environments with many similar distractors, stimuli should be pilot-tested with multiple participants to assess their memorability and recognisability across subjects.

### 2.6 Recording Interviews

As real-world datasets may be susceptible to uncertainty (Shamay-Tsoory & Mendelsohn, 2019; Vallet & van Wassenhove, 2023), collecting high-quality data and capturing the participant’s behavior comprehensively is essential. One useful approach is to gather participants’ subjective experience while participating in the experiment (Kaspar et al., 2014; Mühlinghaus et al., 2024; Susilo & Kitamura, 2005). This can provide valuable context for interpreting data (Lahlou, 2011), help discover motivations or misunderstandings not visible in the behavioral data alone, and capture subjective experiences (Kaspar et al., 2014; Mühlinghaus et al., 2024; Susilo & Kitamura, 2005).

We prepared several unstructured interviews with our participant and documentation of each by video taping with a GoPro 10 for analysis afterwards.

Having participants narrate individual task sessions might provide insights into spatial knowledge acquisition (Viaene et al., 2014).

### 2.7 Effects of Weather and Sunlight

Unlike controlled laboratory settings, outdoor data collection will have to contend with fluctuating environmental variables such as lighting, temperature, and weather. These variables can affect participant performance and equipment functionality (Evans et al., 2012; Hancock and Vasmatzidis, 2003; Salari & Bednarik, 2024) and, therefore, have to be considered during experimental preparation.

We collected our data outdoors in Cyprus during late summer, which came with advantages and challenges regarding weather and sunlight. Many daylight hours, necessary for good video quality of the eye-tracker’s world camera and the GoPro, were beneficial for long data collection days. Additionally, the minimal risk of rain was a practical advantage for using technical equipment in an outdoor setting. However, this period also presented challenges related to intense sun and heat. Direct sunlight can affect video quality due to overexposure and potentially impact eye-tracking accuracy. We successfully dealt with the latter by having the participant wear a sun hat shielding the eye-tracker. The more significant challenge was managing the extreme heat, which required frequent breaks for rest and hydration for the participant and the equipment. During these breaks, we held the overheated devices in the air and removed the batteries to help them to cool down efficiently. Nonetheless, one phone used for GPS data recording overheated and required replacement for the final day of data collection.

We recommend carefully evaluating the weather when collecting data outside. If there is a possibility of rain, we recommend carrying an umbrella. In the case of sunlight, a sun hat can shield sensitive devices, such as the eye-tracker, from direct sunlight (Evans et al., 2012).

Furthermore, it is good to consider how to cool down devices between recording sessions. This might involve scheduling frequent breaks in shaded areas, using portable fans or cooling packs, or avoiding recording during the warmest hours of the day. Finally, a backup plan for equipment failure as a result of weather conditions might be important, including spare devices and a flexible recording schedule.

### 2.8 Equipment and Piloting in the Real World

Environmental variability, timing uncertainties, and the absence of automated controls influence real-world experiments (Stangl et al., 2023; Vallet & van Wassenhove, 2023). Because of these factors, thorough piloting is essential for identifying and resolving potential issues to improve data reliability, participant comfort, and overall study robustness (Niehorster et al., 2025; Salari & Bednarik, 2024).

Before data collection, we thoroughly evaluated the experimental setup. We assessed eye-tracker performance with and without calibration, finding no significant difference and therefore choosing the uncalibrated configuration. We systematically tested the eye-tracker validation under varying lighting conditions and evaluated the effectiveness of a sunhat in mitigating glare. Additionally, we compared head-tracking performance to a compass and identified the needed distance between the two devices to avoid interference. Furthermore, we examined GPS accuracy in the case of an internet connection loss and challenging environments such as narrow streets. Finally, we practiced the complete experimental procedure to ensure a smooth workflow before collecting data in Cyprus. Particular attention should also be given to a realistic assessment of the timing involved in a full recording sequence including all verification and validation procedures to avoid timing issues during the recordings on-site.

While common practice, we recommend not cutting corners but instead carefully piloting each experimental aspect and practicing the different procedures. This should include assessing the functionality of all equipment, determining optimal device positioning, testing GPS accuracy in realistic environments and a realistic timing assessment of a full recording segment.

## 3 Performing the Experiment on Site

In controlled lab experiments, most experimental variables, including the experimental duration, number of trials, and equipment monitoring, can be preprogrammed and require minimal oversight. Adapting such experiments to a real-world setting demands careful consideration and potential adjustments of these variables. In the following, we outline important aspects of performing experiments and strategies for reliably collecting data in the real world, drawing insights from our spatial navigation study.

### 3.1 On Site Logistics

When moving from the virtual to the real world, logistical challenges are essential to consider for reliable data collection. This includes considerations such as power supply or the challenge of transporting the participant to relevant locations, such as the starting location of each exploration session or the task buildings for the pointing tasks. In VR, these aspects can be easily dealt with. For example, participants can be teleported between locations to avoid navigational cues (i.e., Sánchez Pacheco et al., 2025; Schmidt et al., 2023), which is unfortunately impossible in the real world.

Instead, we drove the participant blindfolded to the experimental area and then relied on blindfolded navigation to guide her between locations, at the beginning of the experiment, after each 30-minute session, and for each task location. As guiding the participant without visual input was too time-consuming (see Section 3.7 *Time Management* for details), we used a semi-blindfolded navigation on-site. The participant used her sunhat to obscure visual input other than the ground directly before her feet to help with walking. At the same time, the experimenter guided her on different routes and asked her to spin in circles regularly to lose orientation. While the semi-blindfolded navigation worked relatively well, occasional cues from the ground during the blindfolded navigation required the participant to actively avoid using the ground for navigation during the exploration and task sessions, which might be more challenging to achieve with naïve participants. Other than seeing the ground, the participant additionally experienced other sensory cues, such as sounds and smells, linked to specific locations within the city, as well as internal body cues, including vestibular, proprioceptive, and motor signals, that may have influenced her spatial awareness.

Beyond participant transportation, effective equipment management is crucial. We carried three power banks to charge all necessary equipment during recording breaks. Recharging all devices, including the power banks, upon returning to the hotel presented a logistical puzzle, as we only had two power outlets available. This experience highlighted the importance of anticipating such practical challenges outside a controlled lab environment. Finally, to ensure that we brought all necessary equipment each day, we used a detailed packing list before leaving the hotel.

To minimize participants’ exposure to environmental cues during transitions, we recommend using (semi-)blindfolded navigation, potentially in combination with noise-cancelling headphones or a concurrent cognitive task to reduce attention to surroundings (Barhorst-Cates et al., 2020; Rand et al., 2015). Blindfolding the participant worked well in our study; maintaining this level of unawareness may be more challenging with naïve participants. We therefore recommend minimizing transitions whenever possible. Depending on the setting, transporting participants by car can be an effective method. Additionally, real-world experiments necessitate considering mobile power supply needs and creating a detailed packing list.

### 3.2 Preparing the Equipment

Collecting reliable data in outdoor experiments requires not only a well-calibrated multi-device setup but also a clear and repeatable procedure to synchronize and validate all systems (Franchak & Yu, 2022; Fu et al., 2024; Ladouce et al., 2017). The complexity of coordinating different devices, often with temporal or functional dependencies, start-up times, and data formats, makes a structured preparation phase essential for minimizing errors and ensuring high data quality.

To ensure accurate setup and synchronization of the different devices, we implemented a calibration and validation procedure executed at the beginning and end of every ten-minute recording segment (Sánchez Pacheco et al., 2025; Schmidt et al., 2023). This protocol followed a fixed order to ensure proper device function and synchronization of data streams during the analysis. The procedure began with connecting the RTK GPS system using the NTRIP profile (Weber et al., 2005) and starting GPS recording. Next, we turned on the GoPro, allowing its GPS to stabilize for several minutes, something that is required for reliably recording GPS data. We then tested the eye tracker’s world and eye cameras, followed by calibrating the built-in head tracker with 360° rotations along all three axes, as recommended by Pupil Labs (Baumann & Dierkes, 2023). Once this calibration was complete, we placed and aligned the eye tracker with the wooden sword’s compass and started the recording. Two validation steps followed. First, to later compare and validate the head-tracking (IMU) data offline, we took a picture of the compass’ direction. Second, to assess the eye tracker’s accuracy, the participant put on the glasses and subsequently performed a ten-point validation using a handheld target at a distance of three meters (see Figure 2E), following the procedure created for Pupil Labs Core (Kassner et al., 2014). After these steps, the GoPro recording was started. For organizational purposes, the date, time, and session number were shown to the eye-tracking and GoPro cameras (see Figure 2D). Finally, the participant performed a short serpentine walking pattern for offline synchronization of the GPS and eye-tracking recordings (see Section 3.3 *Temporal Alignment of Data Streams* for details). With all systems recording, the exploration session began.

**Figure 2:**
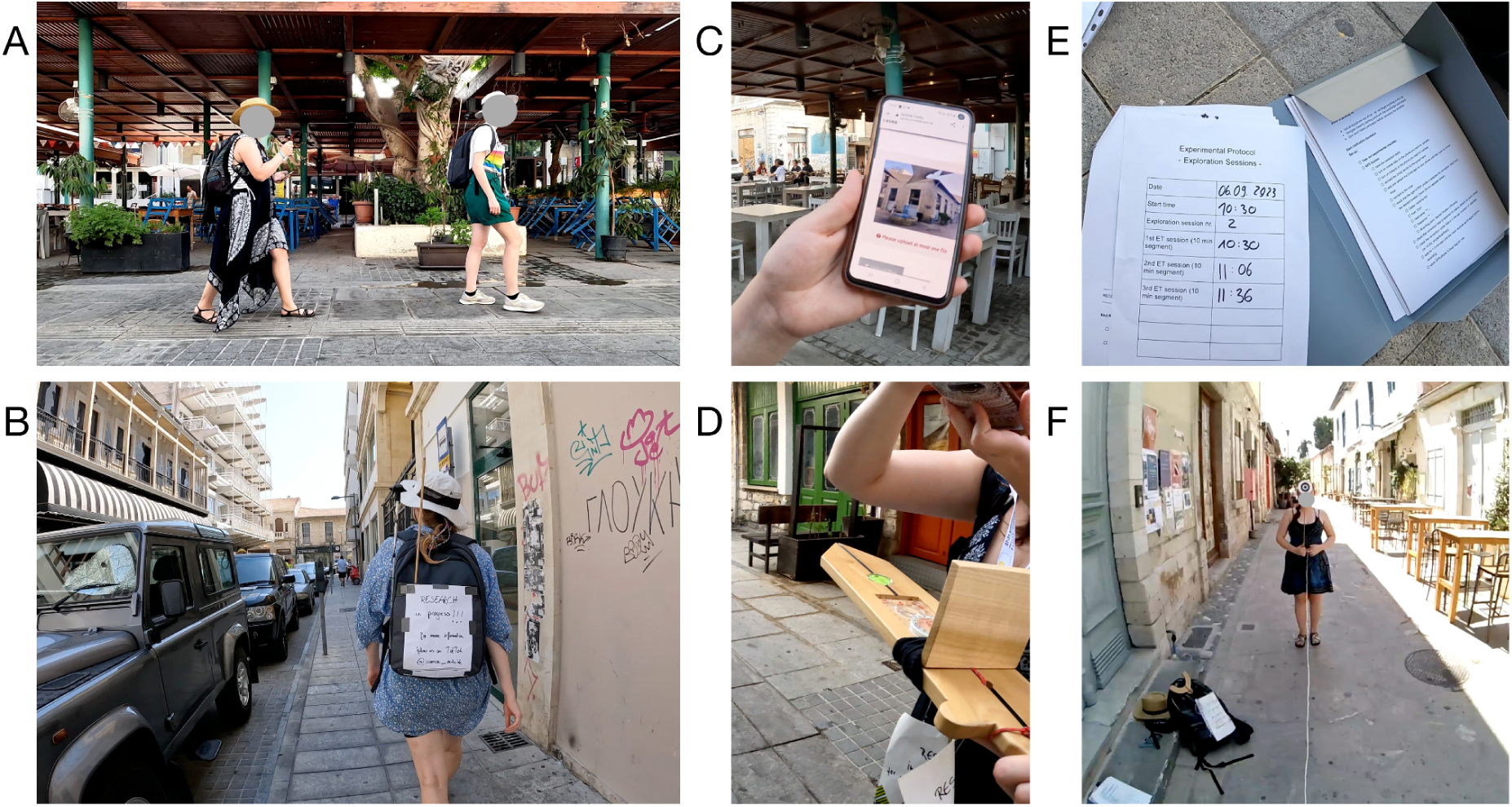
Performing the Experiment. (A) The exploration phase. The participant was freely exploring the city with the experimenter following closely behind her, recording the participant using a GoPro. (B) Snapshot of the exploration phase. (C) For the pointing-to-building task, one of the task buildings is shown to the participant using the experimenter’s phone. (D) For the pointing-to-north task, the participant is pointing towards where she believes north to be. The experimenter is recording her performance by taking a picture of the compass reading. (E) The current session protocol (left) and an empty checklist (right), to help with the equipment setup and later data organization. The session protocol was shown to the GoPro and eye-tracking world camera before the start of each exploration and task session. (F) The participant’s point of view during the eye-tracking calibration. The experimenter was holding a target at ten different locations for the participant to focus on. To ensure a consistent distance, the experimenter and participant spanned a three-meter-long rope between them (the white line in the image).

After each ten-minute exploration segment, the same steps were completed in reverse order: synchronization movements were repeated, the GoPro was stopped, and the accuracy of the eye tracker and head tracker was reassessed. Recordings were then stopped on all devices. If the session marked the end of a 30-minute exploration block, the participant was blindfolded and guided back to the starting point (see Section 3.1 *On Site Logistics* for details). Otherwise, a break was taken to cool down equipment, prevent overheating, and recharge the devices if necessary.

The pointing task followed a similar procedure but excluded device synchronization. The participant was stationary, and precise alignment between GPS and eye-tracking data was not needed. Instead, the exact task locations were manually marked in Google Maps.

Based on our experience, we recommend developing a structured protocol for equipment calibration, validation, and synchronization to ensure high data quality and trust in the recordings. Standardizing this process across sessions can help to guarantee consistency and prevent errors in multi-device setups.

### 3.3 Synchronization of Data Streams

When collecting time-series data from different devices, the temporal alignment of the different data streams is crucial for accurate data analysis (Ladouce et al., 2017). LabStreamingLayer (LSL; Kothe, 2014) is an easy and reliable software solution that aligns different data streams and has been reliably used in various studies (Park et al., 2018; Wang et al., 2023). However, if LSL is not available or possible to use with the recording devices, other options must be sought.

We needed to align the GPS data with the data recorded by the eye tracker. Due to time constraints and the absence of an available LSL integration for our GPS device, we relied on manual synchronization. The approach involved performing a distinct movement pattern that would be identifiable in the GPS and head-tracking data of the eye-tracker. This way, we could roughly align the two data streams offline. After testing several options, we settled on a serpentine walking pattern: the participant walked away from the experimenter and returned, followed by an auditory signal, immediately before and after each ten-minute exploration session. This pattern and sound helped to select the precise recording start and end and enabled us to align the data streams offline by visually inspecting the GPS trajectories and head movements (see Section 4.4 *Temporal Alignment of Data Streams* for details).

For future studies, we recommend using software-based synchronisation, such as LSL whenever possible. This is especially relevant when a high temporal precision is necessary, for example, for EEG (Luck, 2014). A manual synchronization method proves robust and can be recommended as a fall-back solution when combining data streams, e.g. GPS, where the exact temporal precision is less relevant, and might serve as a low-tech control.

### 3.4 Spatial Exploration Phase

With the emergence of untested real-world experimental designs using novel technologies (Ladouce et al., 2017; Shamay-Tsoory & Mendelsohn, 2019), the demand to track technical details and participant behavior may be higher than in controlled lab studies. As a result, recording an exploration phase, during which a participant freely moves with minimal instructions, may require careful planning and continuous monitoring of technical equipment.

During the recording session in our experiment, the participant focused on navigation (see Figure 2A) while periodically checking that the phones recording eye-tracking and GPS data did not overheat. This task’s responsibility lay with the participant, as she was the one physically carrying the phones. The experimenter, following at a one-meter distance (see Figure 2A), managed technical oversight, monitored time using her phone’s timer, recorded participant behavior using a GoPro, and identified suitable break locations for calibration and validation. Additionally, the experimenter followed the participant’s movement using Google Maps to ensure she stayed within the experimental area. The participant was unaware of the exact experimental boundaries, so the experimenter verbally informed her whenever she tried to pass them.

Moving from lab setups to the real world can increase the cognitive load and task responsibilities for the experimenter, which should be accounted for when designing the experiment. To ensure a clean experimental experience, the participant should not be responsible for technical aspects. Instead, one could conduct such experiments with the help of more than one experimenter. Practicing the experimental procedure, like getting familiar with the experimental area before starting, can help ensure smooth execution.

### 3.5 Spatial Navigation Tasks

Conducting experiments, such as spatial navigation tasks, in real-world settings often results in the loss of experimental control (Janssen et al., 2021). Nothing is preprogrammed, so the experimenter must manage task transitions, remember routes, update plans in real time, and adapt to unpredictable surroundings (Lappi, 2015; Vallet & van Wassenhove, 2023). Therefore, conducting navigation tasks in the real world requires careful planning and execution.

Unlike the exploration phase, the experimenter prepared the pointing task setup at each location by herself, while the participant kept her gaze lowered to avoid premature orienting. We used a printed checklist to support the experimenter and ensure smooth setup and execution across trials. Navigation between pointing locations added cognitive demands for the experimenter, as the routes were not pre-planned, requiring her to make navigation decisions in real time to minimize meaningful environmental or navigational cues for the semi-blindfolded participant (see Section 3.1 *On Site Logistics* for details). The pointing task trials were organized in a customized online questionnaire using LimeSurvey, that randomized the task building order, facilitating the data collection and included all experimenter notes during the tasks. At each task location, the participant performed the pointing-to-north task first. The experimenter noted down and also photographed the compass reading (see Figure 2C). For the pointing-to-building task, the pointing targets were displayed to the participant using the experimenter’s phone (see Figure 2B), and responses were logged identically to those in the pointing-to-north trials. After completing the pointing tasks, a map-drawing task was performed in which the participant spent 40 minutes drawing her mental map, including all important buildings and any locations she remembered.

We suggest predefining walking routes between task locations to reduce the need for real-time navigation decisions especially when adapting a semi-blindfolded transport approach. Combining high- and low-tech tools like mobile surveys, printed checklists, and manual compass read outs and pictures may yield methodologically robust real world data collection.

### 3.6 Interviews

We conducted interviews after the first and last exploration sessions, another one during the map-drawing task following the final session, and one to record her reaction after becoming disoriented during one exploration session. They primarily involved the participant reflecting on her decisions, perceptions, and strategies, while the experimenter followed up with guiding questions when necessary.

Our interviews were conducted spontaneously, generally without predefined questions. The experimenter guided the interviews with open-ended prompts (e.g., *“What did you pay attention to?”*), and occasionally added her own observations to help the participant with her reflection. The experimenter also asked follow-up questions when appropriate.

For future studies, we recommend preparing a semi-structured interview guide in advance, but remain flexible to adapt to participants’ behavior. For example, if a participant gets lost, it might be relevant to document this instance immediately. For optimal data collection, researchers should develop guiding questions and a follow-up protocol based on participant responses.

### 3.7 Time Management

A concern for real-world experiments is time management, as data collection may take longer than in controlled lab setups (Vallet & van Wassenhove, 2023). Unforeseen delays and time-consuming procedures can make it difficult to estimate the time needed to record participants adequately.

Despite planning an extra day as a buffer, time management was our biggest difficulty: Recording during the summer in Cyrus introduced the challenge of heat, causing us and the equipment to overheat (see Section 2.7 *Effects of Weather and Sunlight* for details), requiring longer breaks than initially envisioned. On top of that was the overall need for breaks to eat lunch or simply rest our legs. Semi-blindfolded navigation added further time to the exploration phase and experiment (see Section 3.1 *On Site Logistics* for details). Additionally, the need for daylight posed a time constraint, since data recordings after the sun had set was impossible, as it would affect the quality of all cameras, specifically those of the eye tracker. This contributed to us having to cut two exploration sessions short. Furthermore, the equipment setup was very time-consuming, adding, despite prior practice, more than ten minutes before and after each ten-minute recording segment. Finally, many preparations for the experiment were impossible to complete remotely, such as selecting and photographing task buildings or determining the extent of the task area. The time needed to complete them resulted in less time to record data.

For future studies, we strongly recommend scheduling multiple buffer days, thoroughly evaluating the time demands of each step, and optimizing preparation to minimize delays.

## 4 Insights from the Recorded Data

It is important to carefully assess the quality of data collected in real-world environments before data analysis, as real-world environments may introduce different noise sources or rely on novel or rarely tested methodologies. Ideally, such considerations should already be integrated into the preparation and setup phases of the experiment. In the following, we present approaches for validating data collected in real-world settings, with the example of quality assessments from our spatial navigation study.

### 4.1 Eye-Tracking Data

To assess eye-tracking accuracy, we calculated a validation error for each of the ten-point validations. For every target location, we visually identified the relevant time window in the video where the participant fixated on the target. From this window, a one-second stable segment was selected, and five evenly spaced frames were extracted. For each frame, the target’s pixel coordinates were manually annotated using bounding boxes (Tzutalin, n.d.). Simultaneously, corresponding gaze data were averaged to obtain a single gaze point per frame.

The error for each frame was calculated by first obtaining the pixel-based offset between gaze and target location using the Euclidean distance:

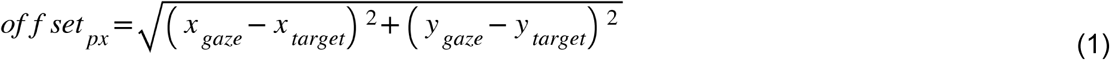

To translate pixel distances into real-world units, pixel offsets were converted to centimeters using the physical size of the targets (a diameter of 12 cm):

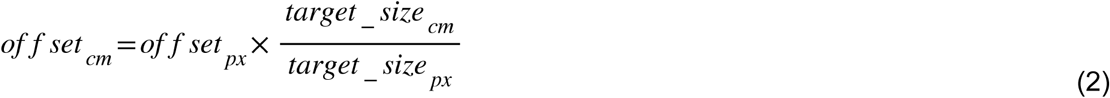

Finally, to express accuracy, offsets in centimeters were transformed to degrees of visual angle, using the known viewing distance of 300 cm:

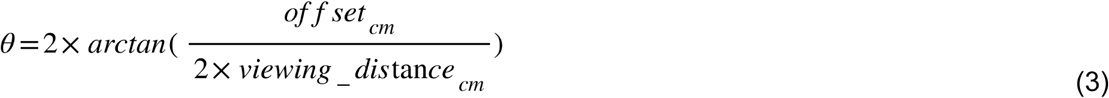

After obtaining five such errors per target, we computed the median error per target and then the median across all ten target positions for each validation procedure. In total, 650 validation points were analyzed across 13 recordings. The results can be seen in Figure 3A. Overall, the start validations had a median angular error of 2.80° (IQR = 1.94 - 3.22). Similarly, the end validations resulted in a median error of 2.65° (IQR = 2.02 - 3.28). A Wilcoxon signed-rank test revealed no differences between start and end validations (W = 30.0, p = 0.31), supporting that after ten minutes of recording, the eye-tracker’s accuracy is constant. Comparing it to Pupil Labs’ reported uncalibrated accuracy of 1.8° (Baumann & Dierkes, 2023), our accuracy is slightly lower than theirs, but given that our real-world setup involved handheld targets and manual identification, we believe this level of accuracy to be very reasonable.

**Figure 3:**
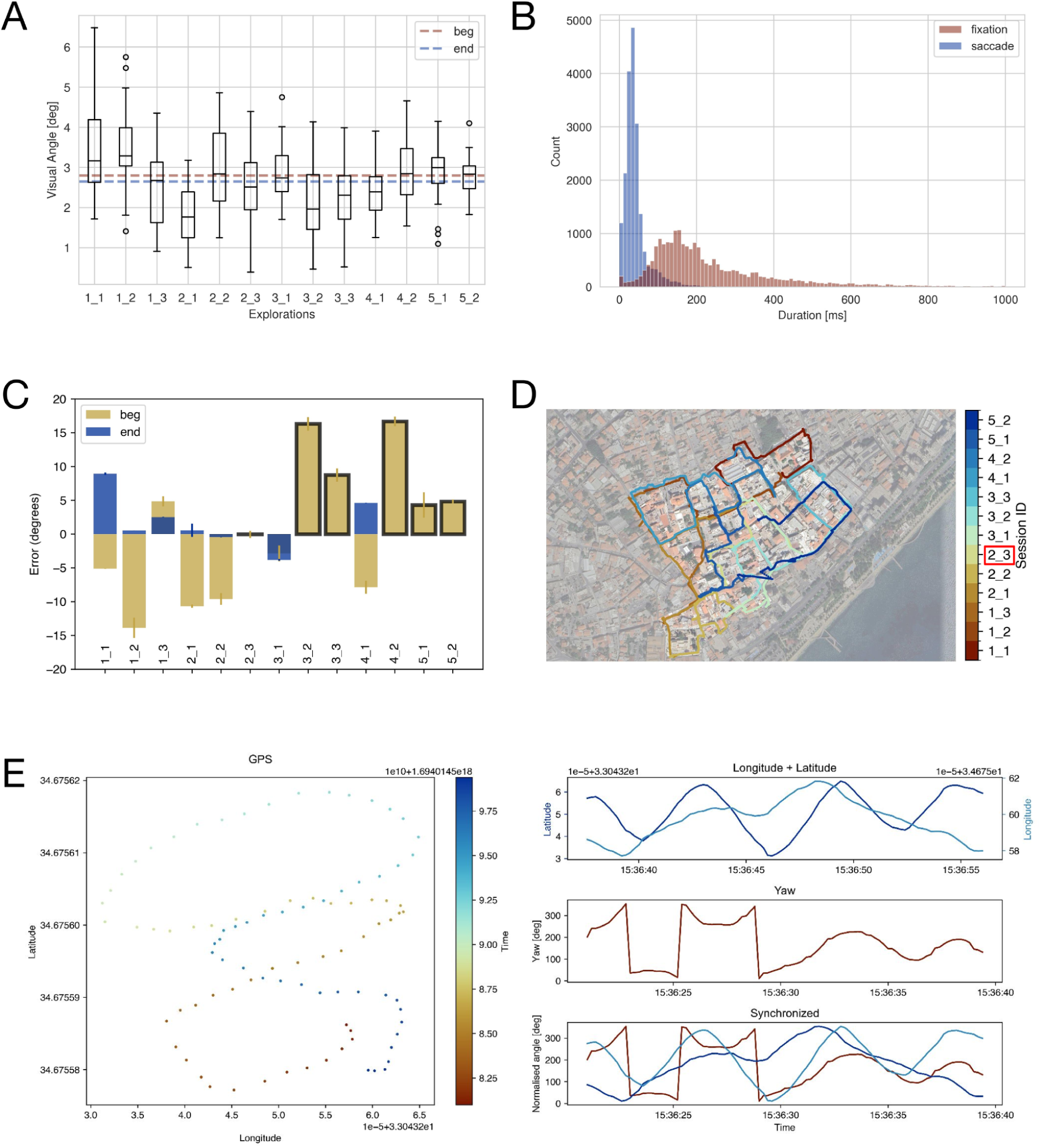
Insights from the Recorded Data. (A) Eye-tracking validation results. Each session is plotted separately, combining all points between the beginning and end. The dashed red line indicates the median error across all start validations, and the blue line indicates the median error for all end validations. (B) The durations of all fixations (red) and saccades (blue) are shown. For each session, we only used the data during the ten-minute exploration. The plot has been cut at 1000 ms for visualization purposes. (C) The difference between the IMU directions and those of the compass. For each session, we show the start (yellow) and end (blue) validations. Black boxes display the sessions with only the start validation. (D) The participant’s position data overlaid over the map of the experimental area. Each session is shown in a different color. The session marked with a red box is the one using the GoPro data. (E) An example of one manual synchronisation. On the left, the GPS movement is shown. On the right, we show the latitude (dark blue), longitude (light blue) from the GPS recording, the yaw movement (red) from the IMU, and the synchronisation after shifting the GPS in time.

Figure 3B additionally shows the fixation and saccade durations across all sessions. The participant had a median fixation duration of 190.55 ms (IQR = 128.27 - 309.43) and a median saccade duration of 34.32 ms (IQR = 24.98 - 46.89).

Device accuracy should always be evaluated relative to task demands: while our measured median accuracy is adequate for large, static targets, finer-grained analyses may require more precise calibration or offline correction. In such situations, mounting calibration targets on fixed surfaces is recommended. The validation targets could also be presented at different distances (Pérez-Edgar et al., 2020), though this step might require additional time investments. Beyond the validation, the event durations of our recorded data are relatively similar to patterns observed in previous research (Dar et al., 2021; Pannasch et al., 2008), supporting the feasibility of collecting reliable eye-tracking data in real-world conditions without significant loss in quality.

### 4.2 Head-Tracking Data

To evaluate the accuracy of the head tracker integrated in the Pupil Labs Neon system (Baumann & Dierkes, 2023; see Figure 1A), we contrasted the head-tracking directions along the yaw component, with those recorded by a compass before and after each recording segment (see Section 3.2 *Preparing the Equipment* for details). The compass values were extracted from the corresponding photos for each validation and converted to cardinal degrees. We then extracted a 0.5-second window (55 samples) from the magnetometer data of the head-tracker to calculate a median heading. For the validation at the beginning of the recording, we used the first 55 samples of the recording. To determine the head tracking orientation for the end validation, we identified the timestamps in the world camera video where the tracker was placed on the sword, and used the first 55 samples afterwards. The head tracker’s directional output (–180° to 180°) was transformed to cardinal degrees and compared to the compass values to calculate errors. The results can be seen in Figure 3C. Of note, due to an internal PupilLabs error, which has been resolved since, some recordings were missing the last 1.5 minutes of the head-tracking data. As a result, we could not evaluate the end validation of several sessions. Overall, we achieved a median error of 0.55° (IQR = −4.15 - 4.83) across all validations. Contrasting the start and end validations, only for sessions where both were present, using a Wilcoxon signed-rank test revealed no significant differences (W = 3.00, p = 0.078), meaning the head-tracker’s accuracy did not change over time.

We recommend that future studies conduct similar validation procedures, especially when the equipment is using a magnetometer and when the head-tracking data is central to the addressed theoretical question. When aligning a head tracker with a compass, the compass’s accuracy can be affected by the presence of other electronic devices, and similarly, the head-tracker’s accuracy can be impacted by the presence of a compass. Therefore, performing validation checks requires the two devices to be placed at a sufficiently large distance from one another.

### 4.3 GPS Data

To process and clean the RTK’s GPS data, we first exclude all manually identified calibration phases (see Section 4.4 *Temporal Alignment of Data Streams* for details), leaving us with data from the end of the start calibration to the beginning of the end calibration. Then, movement speed was calculated based on the distance and time difference between consecutive data points. We first flagged outliers using strict criteria: a quality score of five, fewer than five satellites, or walking speeds exceeding 6 km/h. If applying these criteria would result in more than 30 seconds of consecutive missing data, a more relaxed set of thresholds was used instead: the same quality value of five, fewer than four satellites, or speeds exceeding 7 km/h. Outliers identified by either method were then linearly interpolated. For the session 2_3 where the RTK failed, we used the GoPro’s GPS data instead. Here, outliers were flagged if the movement speed exceeded 6 km/h or if the precision exceeded a threshold of the mean precision plus three standard deviations. As with the RTK’s GPS data, outliers were rejected and linearly interpolated.

Across all sessions, the participant explored the city with an average movement speed of 3.35 km/h ± 1.15, which is a lot slower than the movement speed in VR where participants could move at a speed of up to 18 km/h (Sánchez Pacheco et al., 2025). She covered the entire area, but did not visit many sections more than once. Her exploration pattern is displayed in Figure 3D. Overall, the GPS data aligns well with the street layout, confirming the general reliability of our systems. Only some sections (e.g., the beginning of the recording segment 5.2 in Figure 3D) could not be recovered, resulting in slightly inaccurate data. Interestingly, the GPS data recorded with the GoPro is not visibly different from the data recorded with the RTK, supporting that either option is suitable for such an experiment.

Our results indicate that it is possible to record position data in a real-world experiment accurately. Both the RTK and the GoPro offered stable and accurate position data. While data quality was inaccurate in certain areas and difficult to correct even after thorough cleaning, our methods were generally effective for both RTK and GoPro data. Superb data could be avoided by placing regularly spaced reference markers along the experimental route for data validation and correction, and by scaling the data-collection effort, such as using multiple devices, based on the required level of precision.

### 4.4 Synchronization of Data Streams

To synchronize the GPS data with the IMU signals from the head tracker, we relied on identifiable movement patterns visible in the yaw values recorded by the IMU and in the GPS trajectories (see Figure 3E). For each session, we selected a single time point in both time-streams, either at the beginning or end, based on signal clarity, as the movement pattern was not consistently visible throughout. We manually selected the first timestamp of the calibration movement in the IMU using the video recorded by the eye-tracker’s world camera. This time point was verified through visual inspection of the yaw rotation over time. The corresponding point in the GPS data was determined by plotting the changes in latitude and longitude over time. We then calculated the temporal shift between the IMU and GPS signals using these starting points. This shift was used to adjust the GPS timestamps, aligning the two data streams (see Figure 3E). While visually inspecting the data, we observed that the synchronisations, after which the participant stood still, were the most easily identifiable.

We can recommend our manual synchronisation for future studies with several adjustments for optimal performance. First, performing a distinct movement pattern should be followed by prolonged standing still to improve accuracy, as the absence of movement is easily detectable in the signal. Second, ensuring that the head is aligned with the movement direction improves synchronization precision further. Finally, it is important to note that we could not automate the detection of synchronization points in either dataset. As a result, this synchronization procedure requires manual effort during data analysis.

### 4.5 Task Performance

We assessed spatial knowledge using three tasks: a pointing-to-north task, a pointing-to-buildings task, and a map-drawing task. For the pointing-to-north task, the participant’s location was determined directly from Google Maps markers, and the pointing direction was assessed using the picture of the compass. For the pointing-to-building task, we first obtained coordinates for the participant’s starting location and the building of each task from Google Maps to assess spatial pointing accuracy. The pointing direction was taken from the images of the compass readings and corrected for magnetic declination specific to our location in Cyprus to reference true north. We then calculated the bearing (direction) from the starting location to each task building using the following formula:

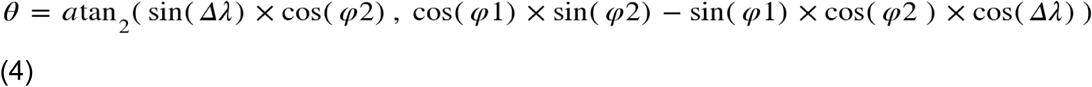

where φ1, λ1 are the latitude and longitude of the starting point and φ2, λ2 the end point, all in radians. To determine pointing accuracy, we computed the difference between the actual and the pointed direction. Figure 4A displays an example of actual versus pointing directions.

**Figure 4:**
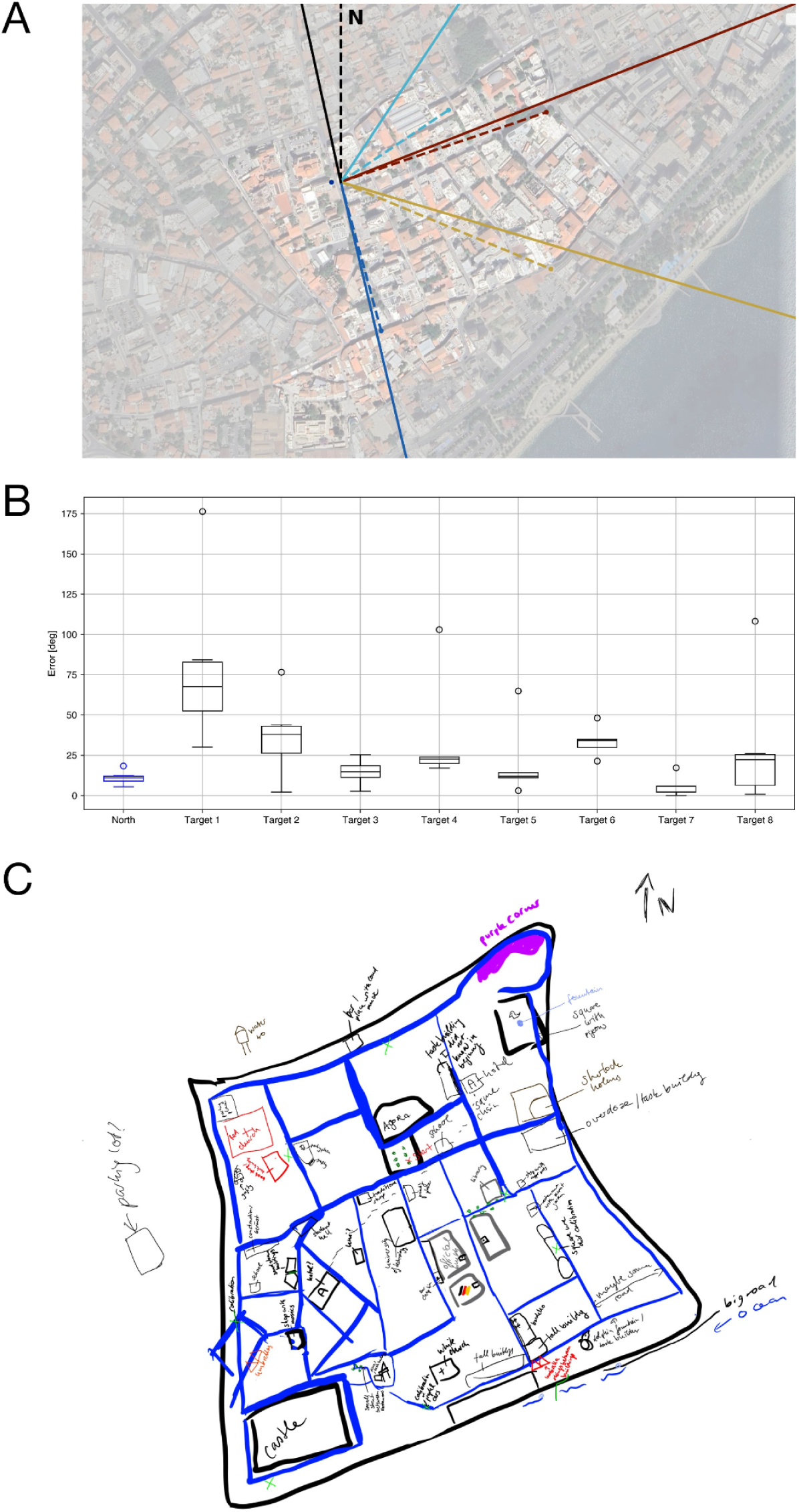
Task Performance. (A) An example of some pointing tasks trials in front of one of the task buildings, indicated by the blue The black lines at the top are the pointing-to-north trial, with the dashed line corresponding to the actual direction of north, and the solid line representing the participant’s response. The remaining colorful lines represent a selected number of pointing-to-building trials. The accurate pointing directions are indicated in dashed lines; the point at the end of each dashed line corresponds to the position of that task building. The solid lines represent the participant’s responses. (B) The participant’s pointing performance for the pointing-to-north (blue) task and the pointing-to-buildings tasks, split up for each task building. (C) The map drawn by the participant. She indicated all streets in blue and the different calibration sessions in green. Each building she remembered is given a short description.

The results of both pointing tasks are presented in Figure 4B. The participant consistently pointed North, achieving an average error of 11.06° ± 4.37. Performance was more variable when pointing to buildings, with an average error of 31.58° ± 33.63. Video recordings during the task revealed that the participant reported several task buildings to be unfamiliar to her, specifically task buildings one and two, which were both only pointed to and not visited during the pointing task. This unfamiliarity reflected in both her performance and confidence levels: task building one had a mean error of 79.54° ± 51.27 and a confidence of zero; task building two had an average error of 36.98° ± 24.61 and a confidence of one. Initially, target building four, also had a low confidence (confidence = 0), but this rose to 10 after the participant completed the pointing task at the location next this target building, accompanied by improved performance (see Figure 4B). Comparing performance and certainty across all buildings using a Spearman’s rank correlation did not reveal a statistically significant association (ρ = −0.619, p = 0.102). Finally, visually comparing the participant’s drawn map and the actual city map revealed many similarities, particularly in building placements and the city’s overall layout (see Figure 4C). However, there were also differences observable. The drawn map was more square-shaped than the real city is, and not all drawn buildings could be recognized or were positioned correctly.

To ensure accurate spatial data collection in real-world experiments, we recommend a multimodal approach using both high- and low-tech options, such as mobile map coordinates, compass directions, and participant reports. We found our handheld pointing device with the integrated compass, paired with mobile surveys and map-drawing tasks, to be an effective tool for assessing spatial knowledge in real-world settings and recommend similar setups for future studies. We further recommend objectively assessing participant familiarity with the task buildings to help with the interpretation of results. Finally, while the map-drawing task provided valuable insights into the participant’s spatial understanding and mental representation of the environment (Coluccia et al., 2007), objectively analyzing such drawings can be challenging. For future studies, we recommend adapting tasks such as the building placement task (Schmidt et al., 2023) for real-world experiments to allow a more objective assessment of survey knowledge.

### 4.6 Interviews

In our study, interview responses reflected both emotional reactions and navigation strategies. Early on, the participant initially divided the experimental area into four sections: left and right of the starting point, and “up,” the area she first walked to, which coincidentally coincided with cardinal north, and “down.” This initial separation was based on her heading direction. She also reflected on her focus during navigation: *“I paid attention to the wrong things… I looked at all the shops, but I won’t recognize them if they don’t have the same assortment.”* Comparing real-world navigation to virtual reality, she noted, *“In VR, you can just bump against things, but in the real world, it might be less fun.”* She additionally commented on the street layout, stating that a lot of streets were very straight and less curvy than expected, and shared how these observations shaped her mental map. When disoriented after a later session, she stated, *“I don’t think I’ve ever been this lost… Yesterday I felt like I knew the city, but now I feel completely lost.”* Finally, while drawing the map, she noted, *“In my head, the city is taller than it is wide,”* and described different layout anchors she had in her mind, something that might be relevant when analysing her behavioral data later on.

The comments and reflections during the interviews helped us understand the participant’s mental representation of space and how spatial judgements were made. The interviews also informed many of the presented best practices and gave insights into different aspects of the experiment. Based on this, we can recommend similar interviews for future studies, as they provide insights into subjective aspects that cannot be assessed otherwise. However, a plan for analyzing the interview data should also be established prior to collecting the data, as similar to the map-drawing task, objectively analyzing such data may be difficult.

## 5 Summary of Best Practices

**Table 1:**
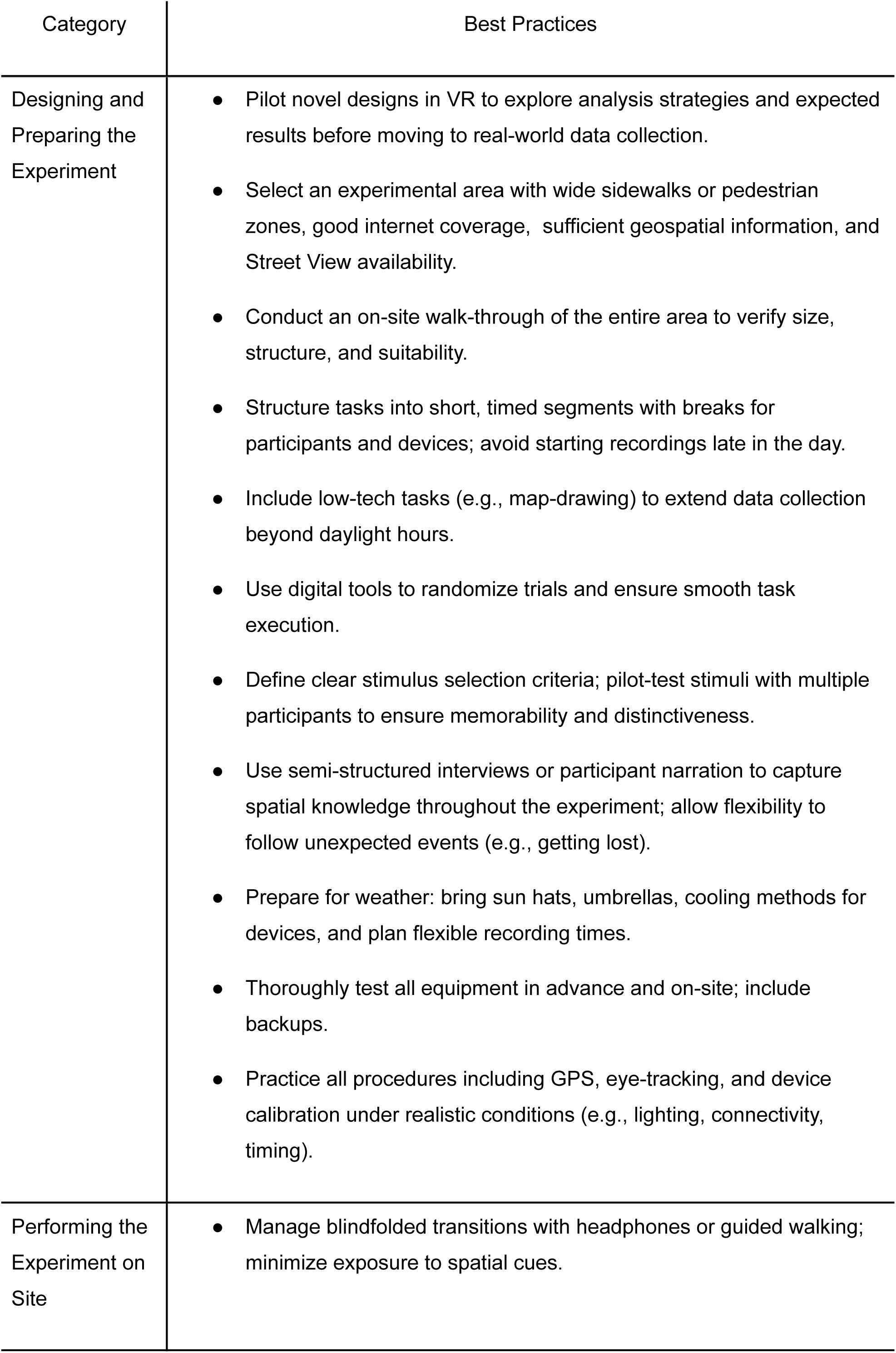
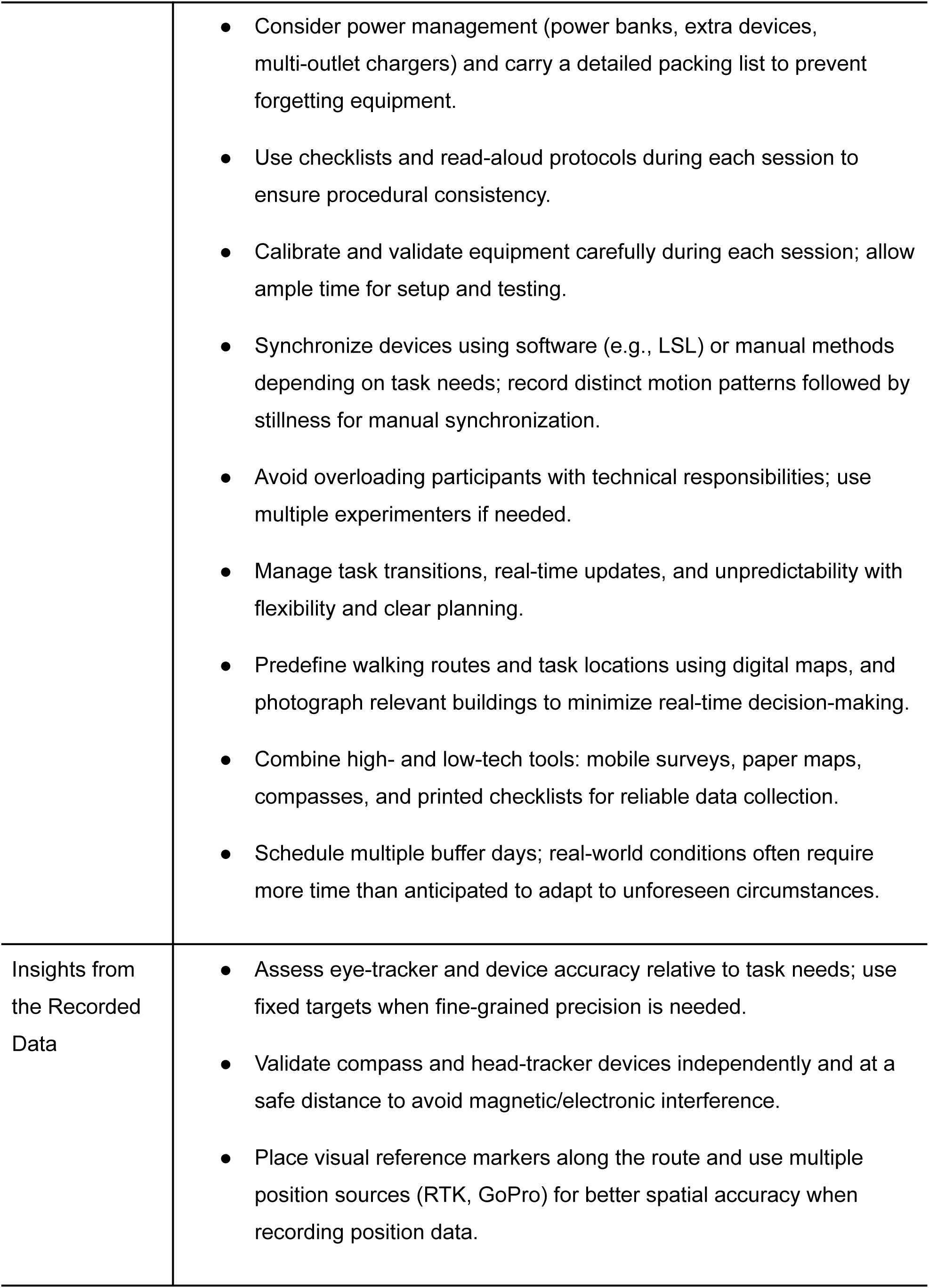
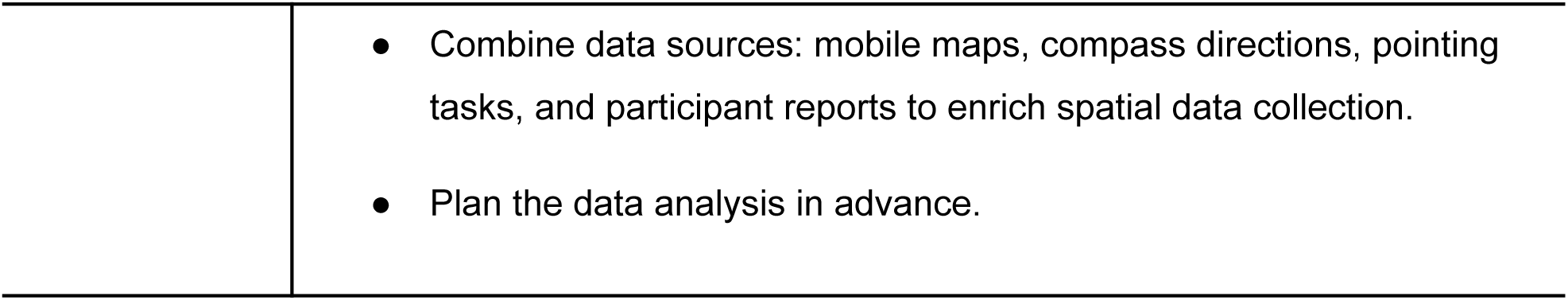
Summary of Best Practices. A table summarising recommendations for designing, preparing, and conducting mobile eye tracking experiments in the real world.

| Category | Best Practices |
| --- | --- |

|  |  |
| --- | --- |
| Designing and Preparing the Experiment | • Pilot novel designs in VR to explore analysis strategies and expected results before moving to real-world data collection.<br>• Select an experimental area with wide sidewalks or pedestrian zones, good internet coverage, sufficient geospatial information, and Street View availability.<br>• Conduct an on-site walk-through of the entire area to verify size, structure, and suitability.<br>• Structure tasks into short, timed segments with breaks for participants and devices; avoid starting recordings late in the day.<br>• Include low-tech tasks (e.g., map-drawing) to extend data collection beyond daylight hours.<br>• Use digital tools to randomize trials and ensure smooth task execution.<br>• Define clear stimulus selection criteria; pilot-test stimuli with multiple participants to ensure memorability and distinctiveness.<br>• Use semi-structured interviews or participant narration to capture spatial knowledge throughout the experiment; allow flexibility to follow unexpected events (e.g., getting lost).<br>• Prepare for weather: bring sun hats, umbrellas, cooling methods for devices, and plan flexible recording times.<br>• Thoroughly test all equipment in advance and on-site; include backups.<br>• Practice all procedures including GPS, eye-tracking, and device calibration under realistic conditions (e.g., lighting, connectivity, timing). |
| Performing the Experiment on Site | • Manage blindfolded transitions with headphones or guided walking; minimize exposure to spatial cues. |
|  | • Consider power management (power banks, extra devices, multi-outlet chargers) and carry a detailed packing list to prevent forgetting equipment.<br>• Use checklists and read-aloud protocols during each session to ensure procedural consistency.<br>• Calibrate and validate equipment carefully during each session; allow ample time for setup and testing.<br>• Synchronize devices using software (e.g., LSL) or manual methods depending on task needs; record distinct motion patterns followed by stillness for manual synchronization.<br>• Avoid overloading participants with technical responsibilities; use multiple experimenters if needed.<br>• Manage task transitions, real-time updates, and unpredictability with flexibility and clear planning.<br>• Predefine walking routes and task locations using digital maps, and photograph relevant buildings to minimize real-time decision-making.<br>• Combine high- and low-tech tools: mobile surveys, paper maps, compasses, and printed checklists for reliable data collection.<br>• Schedule multiple buffer days; real-world conditions often require more time than anticipated to adapt to unforeseen circumstances. |
| Insights from the Recorded Data | • Assess eye-tracker and device accuracy relative to task needs; use fixed targets when fine-grained precision is needed.<br>• Validate compass and head-tracker devices independently and at a safe distance to avoid magnetic/electronic interference.<br>• Place visual reference markers along the route and use multiple position sources (RTK, GoPro) for better spatial accuracy when recording position data. |
|  | • Combine data sources: mobile maps, compass directions, pointing tasks, and participant reports to enrich spatial data collection.<br>• Plan the data analysis in advance. |

## 6 Conclusion

As cognitive science increasingly turns to real-world settings to complement traditional laboratory research, there is an important need to establish best practice advice for robust study protocols that ensure high-quality data collection in naturalistic settings. Our single-subject case study replicated two VR spatial navigation experiments (Sánchez Pacheco et al., 2025; Schmidt et al., 2023) within the urban environment of Limassol, Cyprus. By recording and validating behavioral data, including eye movements, head orientation, and GPS trajectories, we demonstrated the feasibility of performing experiments and collecting high-quality data outside the laboratory. Based on our observations, we offer practical considerations and recommendations for researchers seeking to conduct similar real-world studies. While not intended to be exhaustive, these best practices provide a framework for adapting experimental protocols to complex and uncontrolled environments, depending on specific research goals and experimental designs. As technologies for real-world experimentation continue to advance, these guidelines will need to be updated and adjusted accordingly. Furthermore, although the present study focused on a behavioral experiment, the methods we describe should be compatible with mobile neuroimaging approaches, such as EEG and fNIRS (Gramann, 2024; Janssen et al., 2021; Ladouce et al., 2017; Makeig et al., 2009). Our findings suggest that real-world behavior, such as spatial navigation, can be studied in ways that are meaningfully comparable to VR-based experiments (Walter & Nolte, in preparation). As interest grows in mobile, naturalistic approaches, we view our recommendations as a step toward developing methodological foundations for real-world cognitive neuroscience. Ultimately, we hope this practical guidance will support and inspire researchers to move beyond the laboratory and investigate human behavior in complex and rich real-world environments.

## Acknowledgments

This research was supported by the Applied Vision Association (AVA) through the Tom Troscianko Memorial Award in conjunction with the European Conference on Visual Perception (ECVP) 2023, and the authors would like to thank Manuel Spitschan for supporting their application. The authors thank Paula Vondrlik for her help with preprocessing and preparing the eye-tracking data; Louisa Maubach for assisting with experiment design, piloting, and GPS data analysis; and Milad Rouygari for his contributions to the experimental setup and eye-tracking preprocessing. They are also grateful to Vanessa Franke and Luna Döring for their support with various analysis steps, including validating the eye-tracking data. The authors thank Anna L. Gert for her input on eye-tracker usage and Patrick Unterbrink for his assistance in setting up and troubleshooting the GPS equipment. Finally, the authors thank Tracy Sánchez Pacheco, Aitana Grasso-Cladera, and Vincent Schmidt for their helpful insights and feedback throughout the project.

